# Early events in reovirus infection influence induction of innate immune response

**DOI:** 10.1101/2022.02.25.482062

**Authors:** Andrew T. Abad, Pranav Danthi

## Abstract

Mammalian orthoreovirus (reovirus) is a double-stranded (ds)RNA virus which encapsidates its 10 genome segments within a double-layered viral particle. Reovirus infection triggers an antiviral response in host cells which serves to limit viral replication. This antiviral response is initiated by recognition of the incoming viral genome by host sensors present in the cytoplasm. However, how host sensors gain access to the reovirus genome is unclear as this dsRNA is protected by the viral particle proteins throughout infection. To initiate infection, reovirus particles are endocytosed and the outer viral particle layer is disassembled through the action of host proteases. This disassembly event is required for viral escape into the cytoplasm to begin replication. We show that even after disassembly is complete, endosomal proteases are required to induce an immune response to reovirus. Additionally, counter to dogma, our data demonstrate at least some viral dsRNA genome is exposed and detectable during entry. We hypothesize that some proportion of reovirus particles remain trapped within endosomes, allowing for the breakdown of these particles and release of their genome. We show that rapidly uncoating mutants escape the endosome more rapidly and induce a diminished immune response. Further, we show that particles entering through dynamin-independent pathways evade detection by host sensors. Overall, our data provide new insight into how genomes from entering reovirus particles are detected by host cells.

**IMPORTANCE:** Viruses must infect host cells to replicate, often killing the host cell in the process. However, hosts can activate defenses to limit viral replication and protect the organism. To trigger these host defenses to viral infections, host cells must first recognize they are infected. Mammalian orthoreovirus (reovirus) is a model system used to study host-virus interactions. This study identifies aspects of host and virus biology which determine the capacity of host cells to detect infection. Notably, entry of reovirus into host cells plays a critical role in determining the magnitude of immune response triggered during infection. Mutants of reovirus which can enter cells more rapidly are better at avoiding detection by the host. Additionally, reovirus can enter cells through multiple routes. Entry through some of these routes also helps reovirus evade detection.

## INTRODUCTION

The innate immune response is a critical line of defense against viral infections. This response is characterized by the release of interferon (IFN) which signals for cells to initiate an antiviral response mediated by interferon stimulated genes (ISGs)(1). To initiate this response, host cells must first detect viral infection. Host cells rely on pattern-recognition receptors (PRRs) to identify pathogen-associated molecular patterns (PAMPs) to determine the presence of various pathogens(2). One of the major PAMPs used to identify viral infection is nucleic acid. Nucleic acids not normally found in subcellular compartments of healthy uninfected cells or those that do not resemble host nucleic acids are potent PAMPs(2). Because this detection plays a critical role in whether viral infection is successful, viruses have evolved elaborate strategies to minimize detection of viral nucleic acids(3, 4). Understanding viral strategies to limit detection can inform our approaches to combat viral infection.

Infection with mammalian orthoreovirus (herein reovirus), a double-stranded RNA (dsRNA) virus induces an IFN-based innate immune response that can limit infection(5–15). Among these, the induction of type I IFN following reovirus infection is better understood. Reovirus infection triggers an innate immune response mediated by retinoic acid-inducible gene-I (RIG-I) and melanoma-differentiation associated 5 (MDA5), which are members of the RIG-I like receptor (RLR) family of proteins(16–20). When the RLRs interact with their RNA ligands, they signal through the mitochondrial-antiviral signaling (MAVS) adaptor protein, which functions as a signaling platform for the activation of downstream transcription factors interferon-regulatory factor 3 (IRF3) and nuclear factor κ-light chain enhancer of activated B cells (NFκB)(2). These transcription factors promote the expression of type I IFNs to begin the antiviral response. The immune response to reovirus infection is triggered in a similar way to related, and similarly constructed, dsRNA viruses - rotavirus, bluetongue virus and avian reovirus - making reovirus a model system for understanding the immune response to each of these viruses(21).

To establish infection, reovirus, which is comprised of two concentric protein shells, attaches to carbohydrate and proteinaceous receptors at the plasma membrane and it is internalized into endosomes(5). Within the endosome, reovirus particles undergo a series of disassembly steps catalyzed by acid-dependent proteases. This disassembly leads to the loss of outer capsid proteins and formation of pores within the endosomal membrane allowing for deposition of the inner protein shell (core) into the cytoplasm. The core then becomes transcriptionally active, generating capped, plus-strand RNAs that are translated into viral proteins. Newly synthesized core proteins package plus-strand RNAs to form new viral cores which synthesize the minus-strand RNAs. These newly formed cores can undergo secondary rounds of transcription to amplify the replication process or acquire outer capsid proteins for eventual release from the cell.

While it was initially believed that this elaborate replication strategy described above protects the viral genomic dsRNA (a potent PAMP)(17, 19, 20, 22, 23) throughout all of infection, evidence clearly exists that IFN expression dependent on viral genomic material is induced by reovirus infection. An additional fascinating observation is that it is the genomic material within incoming viral particles that is sensed by the RLRs to trigger the type I IFN response(17, 18, 20, 24, 25). In this study, we sought to determine how the incoming reovirus genome becomes detectable by host cells. We demonstrate that transit through the endosomal compartment and the action of endosomal proteases is required for exposure and detection of the genomic dsRNA from a fraction of incoming particles. Our work shows that reovirus mutants that escape the endosomal pathway more rapidly are less readily detected. Reovirus can use multiple uptake pathways even within the same cell type and our results indicate that transit through different pathways produces a different level of innate immune response. Together, our work more clearly establishes a relationship between virus entry and innate immune activation and defines host and viral determinants that can control the detection of reovirus genomic RNA.

## MATERIALS AND METHODS

### Cells

L929 cells obtained from ATCC were maintained in Eagle’s MEM (Lonza) supplemented to contain 10% fetal bovine serum (FBS) (Sigma-Aldrich), and 2 mM L-glutamine (Invitrogen). Spinner-adapted Murine L929 cells were maintained in Joklik’s MEM (Lonza) supplemented to contain 5% FBS, 2 mM L-glutamine, 100 U/mL penicillin (Invitrogen), 100 μg/mL streptomycin (Invitrogen), and 25 ng/mL amphotericin B (Sigma-Aldrich). Spinner-adapted L929 cells were used for cultivating, purifying and titering viruses. Mouse embryonic fibroblasts (MEFs) were maintained in Dulbecco’s MEM supplemented to 10% FBS, and 2mM L-glutamine.

### Virus propagation and purification

All wild-type and mutant viruses used in this study were propagated and purified as previously described(26). Briefly, plaques isolated from plasmid based reverse genetics were propagated successively in T-25, T75 and T-175 flasks to generate P0, P1 and P2 virus stocks respectively. To generate purified virus, L cells infected with P2 reovirus stocks were lysed by sonication. Virus particles were extracted from the lysates using Vertrel-XF (Dupont). The extracted particles were layered onto 1.2- to 1.4-g/cm^3^ CsCl step gradients. The gradients were then centrifuged at 187,000 × *g* for 4 h at 4°C. Bands corresponding to purified virus particles and top component “empty” particles (~1.36 g/cm^3^)(27) were isolated and dialyzed into virus storage buffer (10 mM Tris-HCl, pH 7.4, 15 mM MgCl_2_, and 150 mM NaCl). Following dialysis, the particle concentration was determined by measuring the optical density of the purified virus stocks at 260 nm (OD_260_) (1 unit at OD_260_ is equal to 2.1 × 10^12^ particles/ml)(27). UV-inactivated virus was generated using a UV cross-linker (CL-1000 UV Crosslinker; UVP). T3D virus was irradiated with short-wave (254-nm) UV on ice at a distance of 10 cm for 1 min at 120 mJ/cm^2^ in a 12 well plate.

### Inhibitors

Dynasore was purchased from Cayman Chemical Company. EIPA and E64d were purchased from Sigma. Inhibitors were reconstituted and stored according to manufacturer’s instructions. The inhibitors were diluted in media to the following final concentrations for experiments: 80 μM dynasore, 50 μM EIPA, 100 μM E64d, 20 mM ammonium chloride, 200 μM ribavirin.

### Generation of ISVPs

Virions were digested with 200 μg/ml TLCK (*N*α-*p*-tosyl-L-lysine chloromethyl ketone)-treated chymotrypsin (Worthington Biochemical) in a total volume of 100 μL for 1 hour at 32°C. After 1 h, the reaction mixtures were incubated for 20 min on ice and quenched by the addition of 1 mM phenylmethylsulfonyl fluoride (Sigma-Aldrich).

### Infections

Confluent monolayers of ATCC L929 or MEF cells were pre-treated with inhibitors as indicated in figure legends for 1 h at 37°C in media. Cells were adsorbed with either phosphate-buffered saline (PBS) or reovirus diluted in PBS at the indicated multiplicity of infection (MOI) at room temperature for 1 h, followed by incubation with medium at 37°C for the indicated time interval. All inhibitors were replaced in medium after the 1 h adsorption period. For exogenous IFN-β treatment, 100 U/mL mouse IFN-β (R&D) was added in medium after the 1 h adsorption period.

### IFN-β ELISA

Supernatants were collected following infections in 12 well plates at 24 h post infection. IFN-β levels were determined using Mouse IFN-β DuoSet ELISA Kit (R&D systems). High-binding ELISA plates (Thermo Scientific) were coated with capture antibody and incubated overnight at room temperature. The plate was washed with PBS containing 0.05% Tween 20 three times. The plate was blocked with PBS containing 1% BSA for at least 1 h at room temperature. Standards were prepared as a two-fold dilution series. Following incubation, the plate was washed as before. Standards and samples were plated in duplicate in coated wells. The plate was then incubated for 2 h at room temperature. The plate was washed and the detection antibody was added at the indicated concentration. The plate was incubated for 2 h at room temperature. Following incubation, the plate was washed as before and Streptavidin-HRP was added at indicated concentration. The plate was incubated for 20 min at room temperature in the dark. The plate was washed and TMB Substrate solution (Thermo Scientific) was added and incubated for 20 min at room temperature in the dark. The reaction was stopped following incubation by addition of stop solution and gentle shaking. Optical density was quantified immediately using plate reader Synergy H1 Hybrid Reader (BioTek). Concentration of samples was determined using a standard curve generated from standard samples.

### Plaque assays

Plaque assays to determine infectivity were performed as previously described with some modifications(28). Briefly, control or heat-treated virus samples were diluted into PBS supplemented with 2 mM MgCl_2_ (PBSMg). L cell monolayers in 6-well plates (Greiner Bio-One) were infected with 100 μL of diluted virus for 1 h at room temperature. Following the viral attachment incubation, the monolayers were overlaid with 4 ml of serum-free medium 199 (Sigma-Aldrich) supplemented with 1% Bacto Agar (BD Biosciences), 10 μg/ml TLCK-treated chymotrypsin (Worthington, Biochemical), 2 mM L-glutamine (Invitrogen), 100 U/ml penicillin (Invitrogen), 100 μg/ml streptomycin (Invitrogen), and 25 ng/ml amphotericin B (Sigma-Aldrich). The infected cells were incubated at 37°C, and plaques were counted 5 d post infection.

### Confocal Microscopy

Monolayers of L929 cells were grown on 8-well chambered slides (Falcon) at 37°C to ~80% confluency. Cells were inoculated with T3D at MOI 10 for 1 h at room temperature with gentle rocking. Following incubation, the inoculum was removed and complete media was added with or without ribavirin (200uM). The slide was incubated for 7 h at 37°C. The slide was then washed with PBS and fixed with 3.7% formaldehyde for 10 min at room temperature. The slide was washed with PBS and permeabilized with PBS containing 0.1% Triton-X100 for 10 min at room temperature. When appropriate, cells were treated with 0.5 U/mL ShortCut RNase III (New England BioSciences) in 1X reaction buffer with MnCl2 for 30 min at 37°C. Untreated samples were incubated in reaction buffer without enzyme. After incubation, cells were washed with PBS three times. Cells were then blocked with PBS containing 2.5% BSA for 30 min at room temperature. The blocking solution was then removed and cells were incubated with J2 antibody (1 μg/mL; Scions) and reovirus antisera for 1 h at room temperature in PBS containing 0.1% BSA and 0.1% TX100. After incubation, cells were washed with PBS containing 0.1% BSA and 0.1% TX100 three times. Cells were then incubated with goat anti-mouse Alexa Fluor 488 (1:1000, Invitrogen), donkey anti-rabbit Alexa Fluor 594 (1:1000, Life Technology), and Alexa Fluor Plus 405 Phalloidin (ThermoFisher) for 1 h at room temperature in the dark. Following incubation, cells were washed with PBS containing 0.1% BSA and 0.1% TX100 three times. Chambers were removed from the slide and the slide was rinsed with diH_2_O. A coverslip was mounted with AquaPoly mounting solution (Polysciences). Confocal images were acquired using a Leica SP8 scanning confocal microscope controlled by Leica LAS-X software. Images were obtained using 63x oil-immersion objective and the White Light Laser and 405nm laser and HyD detectors. Three-dimensional image stacks were acquired by recording sequential sections through the *z*-axis. Images were processed with ImageJ (two-dimensional maximum-intensity projections).

### Plate-based indirect immunofluorescence

Cells in 96-well black walled plates were fixed with ice cold methanol at the indicated time as described in the figure legends. Following fixation, cells were washed twice with PBS and then quickly washed with PBS containing 0.5% Tween 20 (PBS-T). Cells were then incubated with reovirus antisera (1:1000) or 4A3 antibody (1:500) for 1 h at 37°C. Following incubation, cells were blocked with PBS containing 2.5% BSA for 1 h at 37°C. Cells were washed with PBS-T three times and then incubated with DRAQ5 (1:10000), sapphire 700 (1:1000), and LICOR IRDye goat anti-rabbit 800CW (1:1000) or LICOR IRDye goat anti-mouse 800CW (1:1000) for cells stained with reovirus antisera or 4A3 antibodies, respectively. Cells were incubated for 1 h at 37°C and washed three times with PBS-T. Water was added to each well and plates were scanned on a LICOR CLx to obtain fluorescence readings at 700 nm and 800 nm wavelengths. A ratio was obtained by dividing the 800 nm reading by the 700 nm reading and background fluorescence was removed by subtracting mock values.

### RT-qPCR

RNA was extracted from infected cells at various times after infection using an Aurum total RNA minikit (Bio-Rad). For RT-qPCR, 0.5 to 2 g of RNA was reverse transcribed with the high-capacity cDNA reverse transcription (RT) kit (Applied Biosystems) using gene-specific primers. cDNA was subjected to qPCR using SYBR Select mastermix and gene-specific primers with StepOnePlus Real-Time PCR System (Applied Biosystem). Fold increases in gene expression with respect to that of control samples (indicated in each figure legend) were measured using the threshold cycle (∆∆CT) method(29).

### Statistical analyses

The reported values represent the means of three independent biological replicates. The error bars indicate standard deviations (SD). *P* values were calculated using Student’s *t* test (two-tailed; unequal variance assumed) for two groups or one-way ANOVA and two-way ANOVA for more than two groups.

## RESULTS

### Reovirus genome is detectable within infected cells

The reovirus genome is thought to be protected by the viral core throughout infection. Yet evidence from multiple laboratories using different cell types suggest that the genome of incoming viral particles is detected to initiate an antiviral response(17, 18, 20, 24, 25). Since the cell is capable of accessing the viral genome for detection, we hypothesized that the viral genome will be exposed and detectable by a dsRNA specific antibody, J2. We therefore probed infected cells for dsRNA and reovirus proteins using confocal microscopy. Cells were also treated with ribavirin to block viral transcription to ensure that any viral dsRNA signal is coming from the incoming dsRNA rather than any newly synthesized viral RNA. We have previously demonstrated that ribavirin reduces the level of T3D viral RNA in L929 cells by 2-3 orders of magnitude(17). At 7 h following infection we can see the presence of dsRNA puncta within infected cells (Fig. 1A). While some of this signal coincided with signal generated by reovirus capsid specific antisera, a significant portion of this signal was in a distinct subcellular location. To confirm that the J2 signal is legitimately from dsRNA, we treated cells with the dsRNA-specific RNase, RNase III, prior to staining. The dsRNA signal is drastically reduced in RNase III-treated cells suggesting that J2 staining specifically recognizes dsRNA in reovirus infected cells under the conditions used (Fig. 1B). Further, dsRNA signal is not observed in mock infected cells indicating dsRNA seen in infected cells is derived from the incoming viral genome rather than the cell (Fig. 1C).

**Figure 1.**
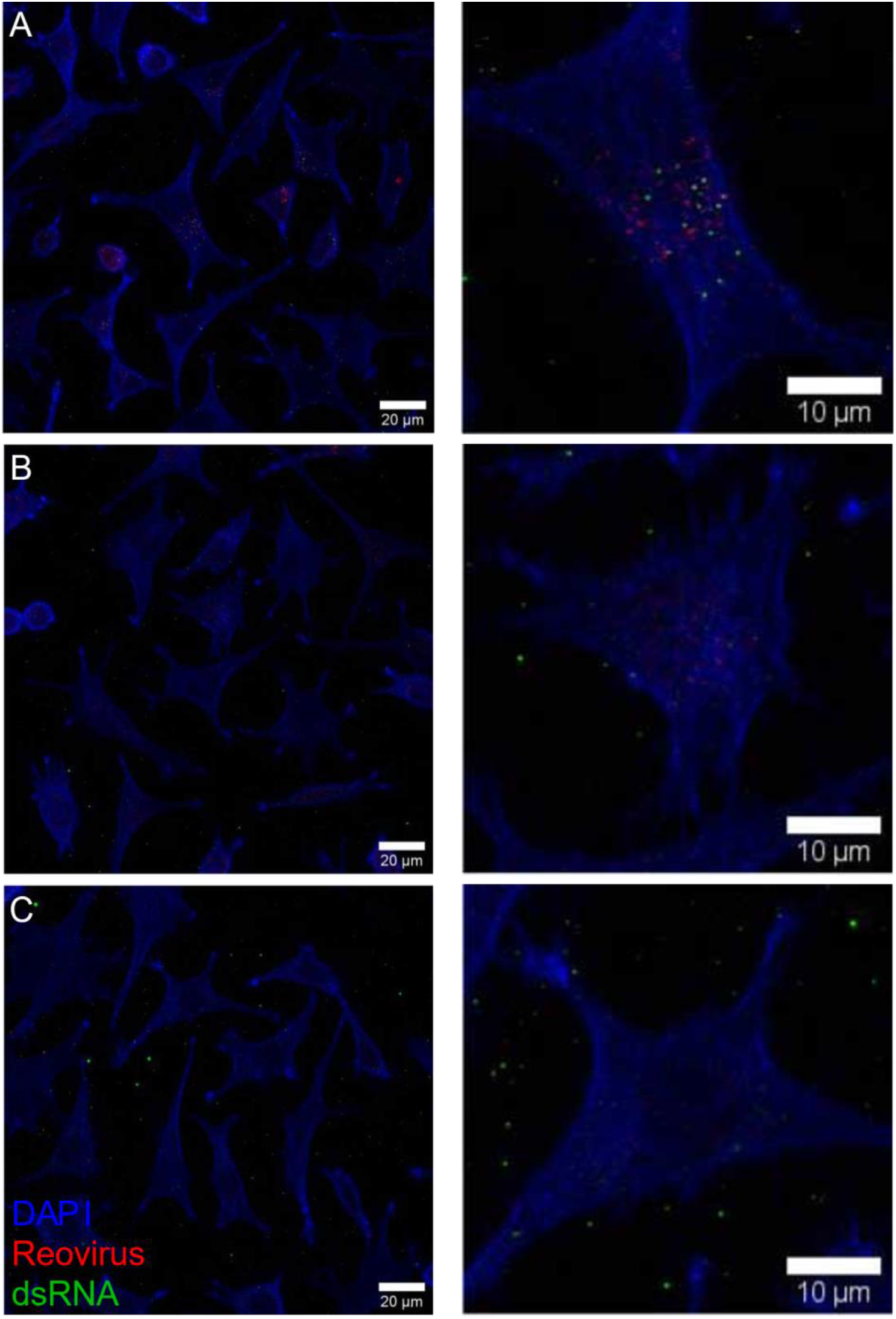
Reovirus genome is detectable within infected cells. L929 cells were infected with T3D at MOI of 10 PFU/cell in the presence of ribavirin and fixed 7 h post infection. Fixed cells were treated without (A) or with RNase III (B) prior to staining for reovirus proteins (red), dsRNA (green) and actin (blue). Uninfected cells are stained for reovirus proteins (red), dsRNA (green) and actin (blue) (C).

### Acid-dependent proteases are required for detection of reovirus infection

To establish infection, reovirus undergoes a series of disassembly steps within endosomes catalyzed by host acid-dependent proteases(5). These disassembly steps lead to the removal of the outer capsid proteins, the formation of pores in the endosomal membrane, and deposition of the viral core into the cytoplasm. Reovirus infection can be blocked by preventing acidification of endosomes using the weak base ammonium chloride (AC) or by specifically blocking cysteine proteases using E64 or its more membrane-permeable derivative, E64d(17, 18, 20, 30–32). Previous work indicates that treatment of cells with AC at the start of infection (0 h post infection) blocks reovirus disassembly and prevents infection(33, 34). Sufficient reovirus disassembly has taken place at 3 and 6 h post infection such that treatment at these times does not influence viral infection(33). To test whether reovirus disassembly is required to generate an innate immune response, we infected L929 cells with T3D and treated cells with AC at 0, 3 and 6 h post infection. At 24 h post infection, we measured cell-associated viral titer. We also measured the type I IFN, IFN-β, secreted in the media of the same cells by ELISA. Consistent with previous work, treatment of infected cells with AC at 0 h post infection drastically reduced viral replication compared to untreated cells (Fig. 2A). The addition of AC at later time points had a lower impact on reducing titer such that treatment at 6 h following infection did not significantly reduce replication. As expected, infection of untreated control L929 cells with reovirus strain T3D triggered a type I IFN response that led to secretion of IFN-β. Preventing disassembly by addition of AC at the start of infection (0 h) significantly reduced IFN-β secretion. Remarkably however, AC treatment at 3 and 6 h post infection also significantly reduced IFN-β secretion (Fig 2B). Additionally, treatment with E64d at 6 h post infection did not impact viral titer but significantly reduced IFN-β secretion (Fig 2A and 2B). These data indicate that acidification of endosomes, and the action of cysteine proteases is necessary for induction of an immune response to reovirus infection. Further, these data show that reovirus replication can be decoupled from an immune response as late treatment (3 and 6 h post infection) with AC or E64d reduces the immune response but has minimal effect on viral replication. Control experiments using cells overexpressing RIG-I, which upregulate IFN-β expression, indicated that these inhibitors do not directly affect signaling via this pathway (data not shown).

**Figure 2.**
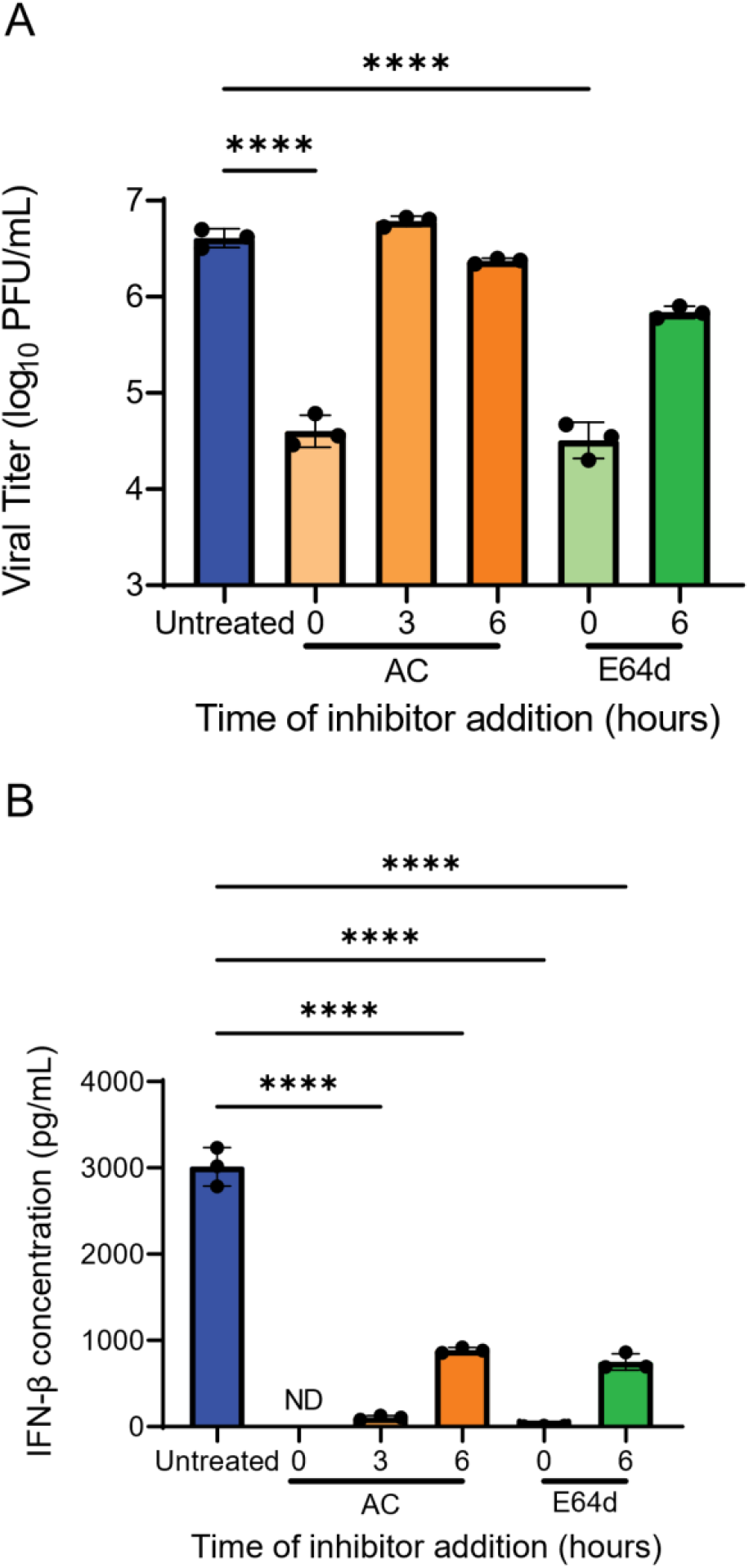
Reovirus replication and induced innate immune response can be uncoupled. L929 cells were infected with T3D at MOI of 10 PFU/cell and treated with 20mM ammonium chloride (AC) or 100μM E64d at different times post infection. At 24 h post infection, titer was measured from cells by plaque assay (A) and secreted interferon-β (IFN-β) was measured from the supernatant by ELISA (B). The data are presented as means and SD of three biological replicates compared to untreated using one-way ANOVA with Dunnett’s multiple comparisons test (**** = p<0.0001)

During entry into host cells, reovirus virions disassemble by the action of cysteine proteases to generate an intermediate known as infectious subvirion particle (ISVP)(5). ISVPs can also be generated *in vitro*. As *in vitro* generated ISVPs are partially disassembled, these particles bypass the requirement for endosomal proteases and can enter the cell earlier in the endosomal sorting pathway or directly at the plasma membrane(35–37). We hypothesize ISVPs evade immune detection by entering the cell without transiting through the degradative environment of endosomes. To test this hypothesis, we compared IFN-β secretion in response to virion or ISVP infection. To ensure that differences observed are not related to differences in the attachment efficiency of these particles, we established conditions under which an equivalent number of virions and ISVPs were attached to the cell (as measured by RT-qPCR for genomic RNA). We found that under these conditions, in comparison to virions, ISVPs induced significantly less IFN-β at 24 h post infection compared to virions (Fig. 3A). *In vitro* generated ISVPs are more infectious than virions in many cell types. It is thought the difference in infectivity between ISVPs and virions is related to inefficiencies in disassembly due to a limited activity of cysteine protease in those cell types(38). Because we found that ISVPs do not induce a significant IFN-β response, it is also possible that the greater infectivity of ISVPs is also related to the absence of the suppressive effects of IFN-β. To test this idea, we infected L929 cells with virions or ISVPs and treated cells with exogenously added IFN-β to a level that is roughly that is produced from virion infected cells. Using indirect immunofluorescence, we observed that ISVPs were more infectious compared to virions but IFN-β treatment reduced ISVP infectivity to levels observed for virions (Fig. 3B). These data indicate that improved ISVP infectivity is at least partially due to lower induction of an innate immune response by entering the cell earlier in the uptake pathway.

**Figure 3.**
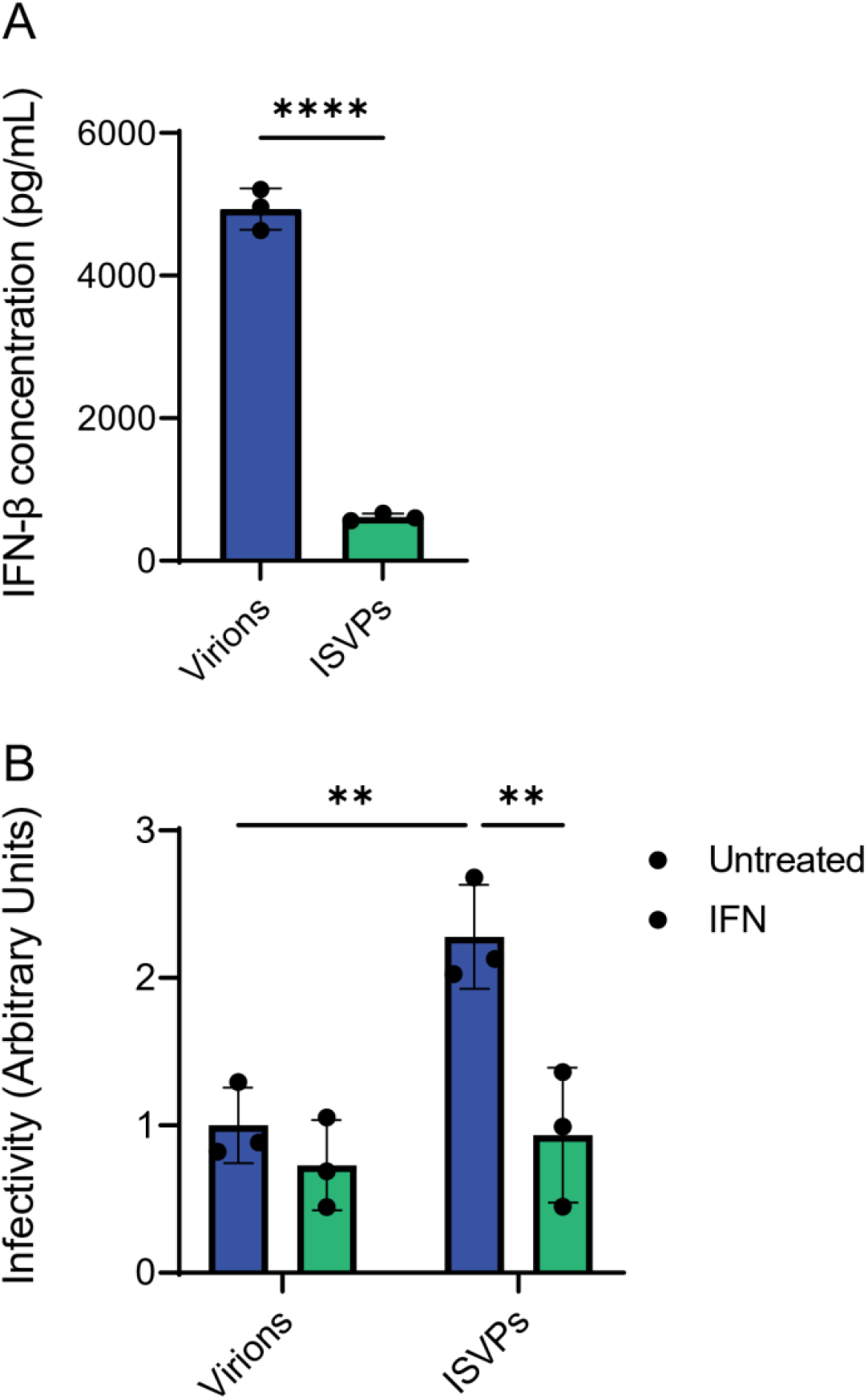
ISVPs evade immune detection through early endosomal escape. L929 cells were infected with T3D virions or ISVPs at 1000 particles/cell. At 24 h post infection, secreted interferon-β (IFN-β) was measured from the supernatant by ELISA (A). The data are presented as means and SD of three biological replicates compared using student *t-*test (**** = p<0.0001). L929 cells were infected with T3D virions or ISVPs at 1000 particles/cell and treated with 100U/mL mouse IFN-β. At 24 h post infection, cells were fixed and infectivity was measured by indirect immunofluorescence for reovirus proteins (B). These data are presented as means and SD of three biological replicates compared using one-way ANOVA with Sidak’s multiple comparison test (** = p<0.01).

### Rapid disassembly of reovirus particles allows for evasion of detection

ISVPs undergo conformational changes that result in formation of ISVP*s. ISVP* formation results in the release of viral pore forming peptides that help the genome-containing core particle to escape the endosomes. Reovirus mutants that undergo ISVP-to-ISVP* conversion at different rates have been identified(39–41). The reassortant T1L/T3DM2 transitions from ISVPs to ISVP*s more efficiently than T1L(39). Conversely, the mutant T1L/T3DM2 D371A transitions to ISVP*s less efficiently compared to the parental T1L/T3DM2 and at a rate more similar to T1L(39). We hypothesize that rapid ISVP* formation decreases the time a particle spends in the endocytic compartment, therefore evading the innate immune response, similar to ISVPs. To test this hypothesis, we first sought to confirm that the *in vitro* observations for differences in the ISVP-to-ISVP* efficiency of these viruses also were recapitulated in infected cells. For these experiments, we stained L929 cells infected with ISVPs of each of these viruses with a conformation-specific monoclonal antibody 4A3. This antibody specifically recognizes the μ1 protein in the ISVP* conformation(42). 4A3 staining therefore allows us to determine the extent to which ISVP* are generated in cells at a given time point. To this end, we adsorbed equal numbers of *in vitro* generated ISVPs of T1L, T1L/T3DM2, or T1L/T3DM2D371A to cells at 4°C. We then measured relative 4A3 staining using indirect immunofluorescence early during infection. Consistent with expectation, no staining was observed at 0 h (data not shown). At 2 h post infection, T1L/T3DM2 showed significantly greater 4A3 staining compared to either T1L or T1L/T3DM2 D371A (Fig. 4A). Thus, these viruses display the same type of difference in ISVP-to-ISVP* conversion in cells as has been reported *in vitro*(39). Additionally, we anticipated efficient disassembly of the particle would allow for increased endosomal escape and core delivery. Therefore, we measured viral transcripts at 2 h post infection as a readout for core delivery and activation. T1L/T3DM2 produced more viral transcripts relative to either T1L or T1L/T3DM2 D371A, indicating increased core activation (Fig. 4B). These data demonstrate that T1L/T3DM2 escapes endosomes more efficiently than slower disassembling viruses.

**Figure 4.**
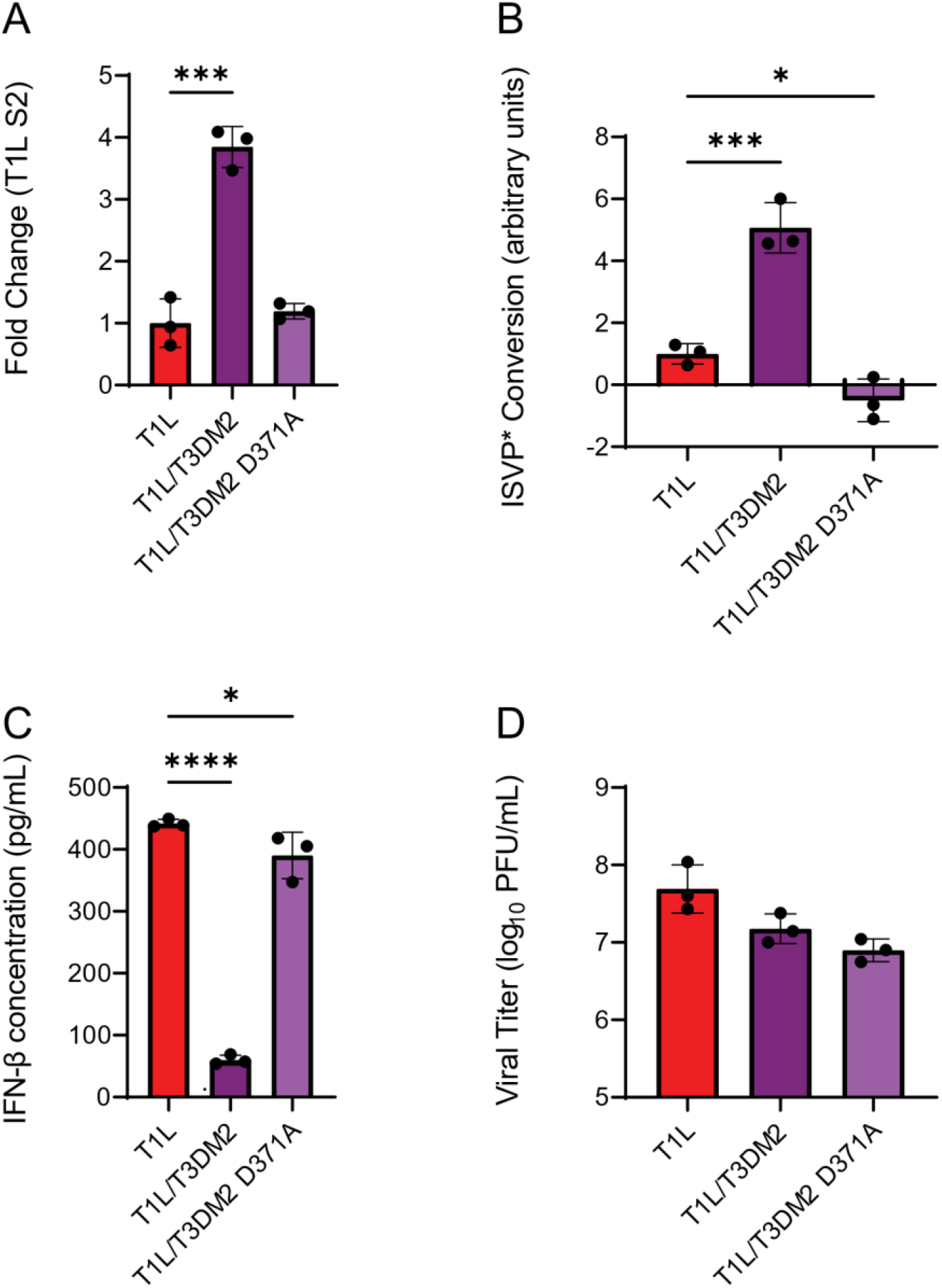
Efficient disassembly of reovirus contributes to immune evasion. L929 cells were infected with T1L, T1L/T3DM2, or T1L/T3DM2 D371A at MOI of 10 PFU/cell and T1L S2 transcripts were measured by RT-qPCR at 7 h post infection (A). L929 cells were infected with T1L, T1L/T3DM2, or T1L/T3DM2 D371A at MOI of 10 PFU/cell in the presence of 100μM ribavirin. ISVP* conversion was measured 2 h post infection by indirect immunofluorescence using the conformationally-specific mAb, 4A3 (B). L929 cells were infected with T1L, T1L/T3DM2, or T1L/T3DM2 D371A at MOI of 10 PFU/cell and secreted interferon-β (IFN-β) was measured from the supernatant by ELISA (C) and viral titer was measured by plaque assay (D). Dotted line indicates limit of detection for ELISA. The data are presented as means and SD of three biological replicates compared using one-way ANOVA with Dunnett’s correction (* = p<0.05; *** = p<0.001; **** = p<0.0001).

Next, we wanted to evaluate how disassembly efficiency of reovirus impacted the immune response. Therefore, we infected cells with each virus and measured IFN-β secretion and viral titer at 24 h post infection. T1L triggered an IFN-β response but T1L/T3DM2 induced significantly less IFN-β during infection (Fig. 4C). In comparison, T1L/T3DM2 D371A induced IFN-β to similar levels to wild-type T1L. Viral titers were similar for all viruses following infection, indicating differences in immune response were not due to differences in viral replication (Fig. 4D). Together, these data indicate differences in rate of ISVP* conversion influences IFN-β induction. Faster disassembling viruses likely spend less time in the endosome, which allows for greater evasion of detection by host cells.

### Mode of entry affects immune response to reovirus infection

Reovirus is capable of using different endocytosis pathways to enter cells to initiate infection(43). While some endocytosis pathways are cell-type specific, in other cases, more than one endocytic pathway can be used in the same cell type. Consistent with this, we found that neither dynasore, which blocks dynamin-dependent endocytosis(44) or 5-(N-Ethyl-N-isopropyl)amiloride (EIPA), which blocks macropinocytosis(45, 46) prevented efficient reovirus replication in MEFs. This result suggests that cell entry in MEFs is not obligately dependent on one of these endocytic pathways (Fig. 5A and 5D). This observation presented an opportunity to test if entry through a specific endocytic pathway impacted the induction of the innate immune response. After confirming that dynasore and EIPA do not impact RIG-I signaling in RIG-I overexpressing cells (data not shown), we evaluated the effect of these inhibitors on the innate immune response to reovirus infection. We pre-treated MEFs with the inhibitor for 1 h prior to infection then allowed infection to proceed for 24 h in the presence of the inhibitors prior to measurement of IFN-β secreted into the supernatant by ELISA. In comparison to vehicle treated control, treatment with dynasore significantly reduced IFN-β secretion in response to reovirus infection (Fig. 5B). These data suggest that the dynamin-dependent modes of endocytosis (dynasore-sensitive) are important for allowing cells to detect reovirus infection. In contrast to these results, EIPA treatment significantly increased IFN-β secretion following reovirus infection compared to vehicle treatment (Fig 5E). Thus, viral particles are better detected when EIPA-dependent uptake is blocked. To determine if changes in IFN-β secretion due to dynasore or EIPA correlated with a more rapid escape from the endosomes, we measured the accumulation of viral transcripts early in infection. In the presence of dynasore, reovirus particles produced significantly more transcripts at 2 h post infection compared to vehicle-treated controls (Fig. 5C). Interestingly, EIPA treatment also increased early viral transcripts (Fig. 5F). These data indicate that the type of uptake pathway can influence the rate at which reovirus launches infection. For this current study, we did not further examine the basis for this observation. Importantly, of greater relevance to this current work, our data suggest that endosomal pathway has a more dominant effect on controlling the innate immune response than the efficiency of escape from the endosomes.

**Figure 5.**
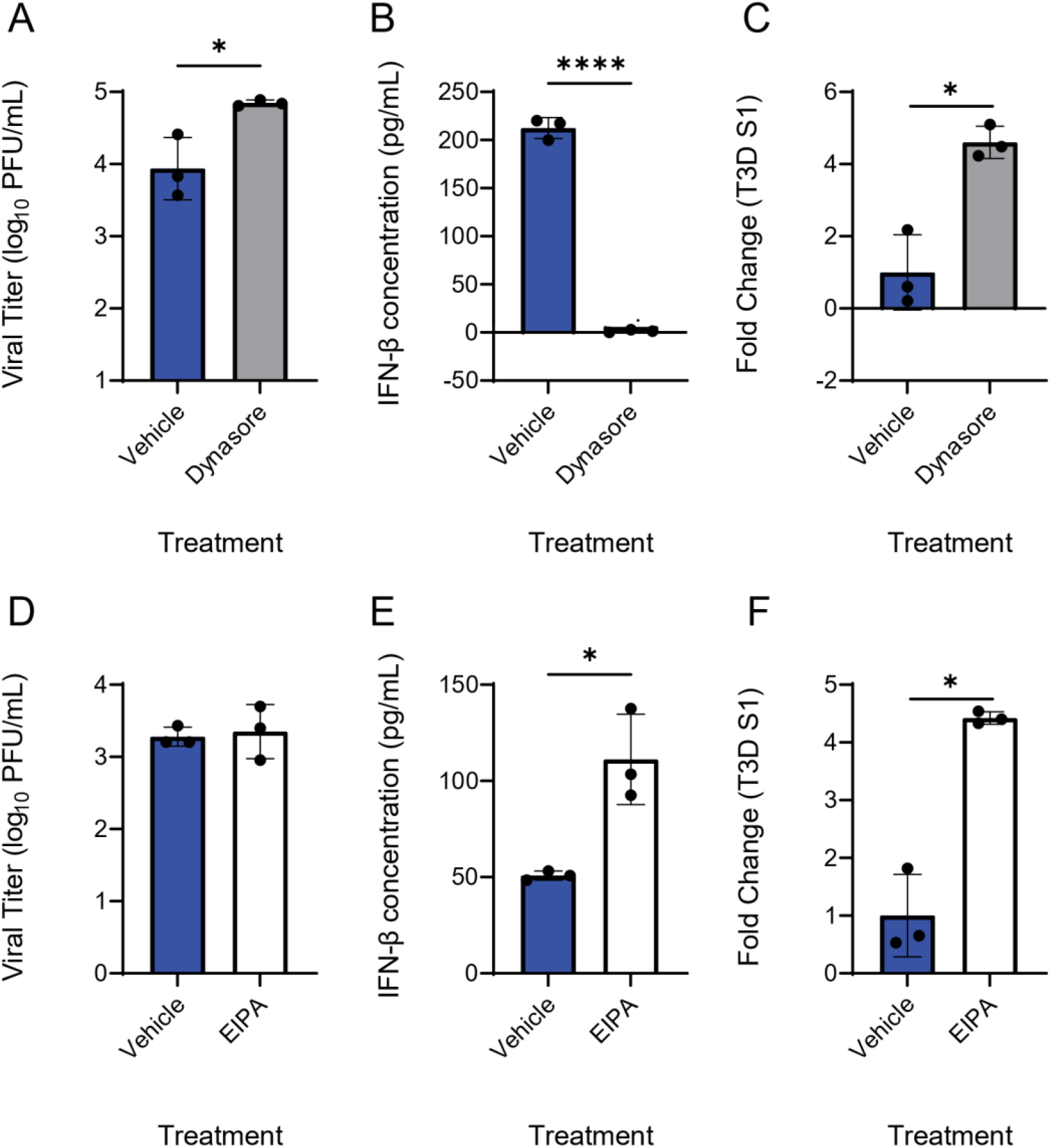
Uptake pathway of reovirus affects innate immune response. MEFs were pre-treated with DMSO or 80μM dynasore for 1 h at 37°C (A-C) or with MeOH or 50μM EIPA (D-E). Cells were then infected with T3D at MOI of 10 PFU/cell in the presence of inhibitor or vehicle control. At 24 h post infection, viral titer was measured by plaque assay (A and D) and secreted interferon-β (IFN-β) was measured from the supernatant by ELISA (B and E). T3D S1 transcripts were measured at 7 h post infection by RT-qPCR (C and F). Dotted line indicates limit of detection for ELISA. The data are presented as means and SD of three biological replicates compared using student *t*-test (* = p<0.05; ** = p<0.01; **** = p<0.0001).

### Uptake pathway of destabilized reovirus particles determines innate immune response

UV-treated reovirus particles bind cells, are disassembled, but fail to activate gene expression efficiently(17, 22, 24, 25). UV-treated particles induce an increased immune response due to destabilization of the reovirus cores and consequently more facile release or exposure of genomic dsRNA(17, 22). To determine whether the detection of genomes from these destabilized particles would no longer be affected by the uptake pathway, MEFs pre-treated with dynasore were inoculated with 1000 particles/cell of T3D or UV-inactivated T3D (T3D-UV). IFN-β was measured from supernatants at 24 h post infection. T3D-UV induced significantly more IFN-β compared to unirradiated T3D (Fig. 6), consistent with previous reports(17, 22, 24, 25). Interestingly, dynasore treatment significantly reduced IFN-β secretion induced by both T3D and T3D-UV. These data suggest that the uptake pathway of reovirus can significantly alter the capacity of host cells to detect incoming particles. Overall, early entry events of reovirus particles dictate how efficiently host cells can sense infection.

**Figure 6.**
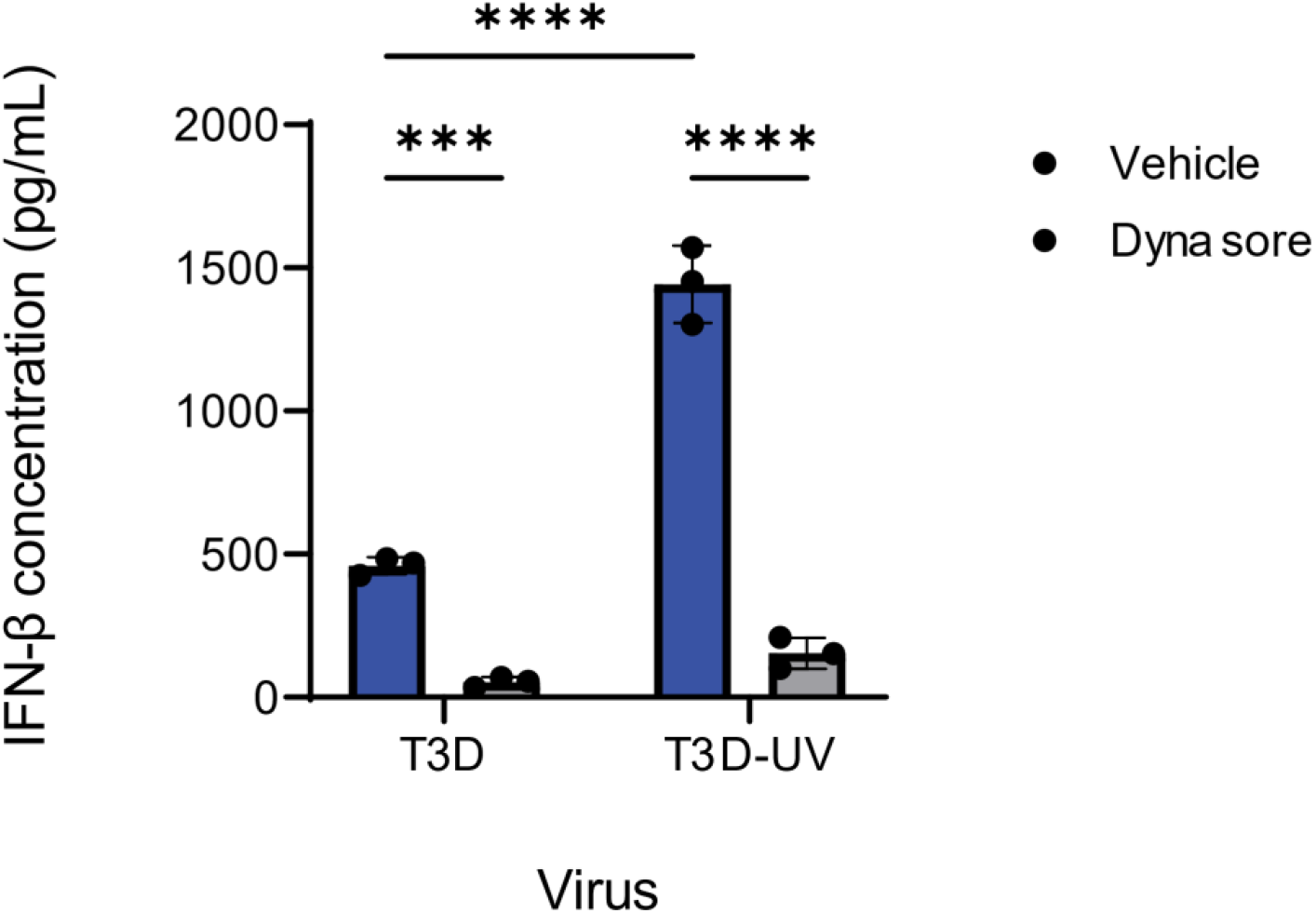
Destabilized reovirus particles are detected by dynamin-dependent uptake pathways. MEFs were pre-treated with DMSO or 80μM dynasore for 1 h at 37°C. Cells were then adsorbed with T3D or UV-inactivated T3D (T3D-UV) at MOI of 1000 particles/cell. At 24 h post adsorption, secreted interferon-β (IFN) was measured from the supernatant by ELISA.

## DISCUSSION

The reovirus genome is thought to be protected by the viral core throughout all of infection(47, 48) yet the incoming genome is sensed to trigger an innate immune response(6–15). In this study, we sought to determine how the incoming viral genome is detected to initiate an immune response during reovirus infection. Contrary to dogma, we demonstrate that the dsRNA genome from incoming viral particles is not fully shielded by the core proteins but is detectable. We show that detection of the genome of incoming particles is dependent on the action of endosomal, acid-dependent proteases, the kinetics with which the virus completes uncoating events to escape the endosome. We also find that efficiency of reovirus detection varies with the specific endocytic pathways used for entry. Together this work provides new insight into how entry events control the innate immune response following infection.

Entry of reovirus virions is an inefficient process with some proportion of particles remaining associated with endosomes at least as late as 4 h post infection(49). It is possible that inefficient reovirus entry contributes to the high particle-to-pfu ratio of ~100:1 for reovirus. If so, numerous non-infectious particles could be broken down when transiting through the endosome to allow for immune detection while the infectious particles, that escape the endosome, contribute to viral replication. The dsRNA genome released from degraded particle is detected by the cytoplasmic RIG-I-like receptors (RLRs)(16–20). TLR3, which functions in the endosome, is dispensable for clearing reovirus infection and for IFN production *in vivo*(50, 51) and in cell culture(17, 20, 38, 52). Therefore, the genome of some proportion of incoming viral particles must reach the cytoplasm to be sensed by the RLRs. One possibility is that some reovirus particles are broken down during disassembly in the same endosome as functional particles such that the released genome enters the cytoplasm simultaneously with functional cores. This possibility is feasible considering multiple reovirus particles can be present within the same endosomal compartment(53). Another possibility for detection is that the viral genome is released within the endosome and transferred into the cytoplasm by a host protein. The endosomal dsRNA transporter, SIDT2, has been implicated in the immune response to other viruses(54). dsRNA generated during infection by herpes simplex virus 1 (HSV-1) or encephalomyocarditis virus (EMCV) could be taken up extracellularly by bystander cells and transferred into the cytoplasm by SIDT2 for immune sensing. We anticipate a similar phenomenon may occur during infection by reovirus for endosomal released genomic RNA.

ISVPs bypass the requirement for these endosomal compartments allowing these particles to enter either directly at the plasma membrane or early within the endosomal sorting pathway(35–37). Since cathepsin activity is required even late in infection to trigger an immune response (Fig. 2), the ability of ISVPs to bypass the cathepsin-containing degradative environment likely contributes to this reduced immune response. When reovirus initiates infections in the gut, it is thought that ISVPs formed extracellularly by the action of luminal proteases are the initiating particle. Based on evidence that ISVPs induce a lower level innate immune response, it is hypothesized that ISVP formation is a viral strategy to evade the innate immune response. Our data supports these observations by showing the increased infectivity of ISVPs is partially due to the reduced IFN response generated during infection (Fig. 3). The ability of ISVPs to avoid detection by initially infected intestinal cells is likely beneficial. Evasion of the immune response could allow the virus to establish infection and replicate to high enough levels before disseminating to other tissue where infection likely produces a robust innate immune response(38).

Following ISVP formation, penetration of endosomal membranes is facilitated by μ1 (encoded by the M2 gene) and changes in this protein can alter the rate of disassembly of the particle(39). We demonstrate that more rapid ISVP-to-ISVP* conversion leads to a decreased innate immune response during infection (Fig. 4). Similarly to initiating infection with ISVPs, we suspect these faster uncoating particles evade the immune response by escaping endosomes more efficiently. In newborn mice, T1L/T3DM2 replicates to higher levels and has a greater myocarditic potential than T1L(55). Our observation that T1L/T3DM2 induces a lower level of type I IFN than T1L is consistent with observations that myocarditic reoviruses induce lower levels of IFN during infection(9). This work is also consistent with evidence that M2 mutants deficient in membrane penetration activate IRF-3 to lower levels compared to wildtype virus(56). Decreased membrane penetration might suggest decreased endosomal escape of these particles allowing for increased immune detection. However, stability of the particle and its potential effect on releasing the genome was not assessed in the context of these mutations. It remains possible that these mutations affect the ability of the cell to break down the particle to gain access to the viral genome, independent of μ1 membrane penetration efficiency.

We show that altering the mode of cellular uptake affects the immune response to virion infection (Fig. 5). We found that skewing virion entry towards dynamin-dependent modes led to an increased immune response indicating these pathways likely contribute to sensing reovirus. Importantly, we found that the uptake pathway is important regardless of the efficiency with which the virus transits the endosomal compartment. How these pathways contribute to detecting reovirus infection is not clear. The composition of endosomes reached when reovirus initiates infection using different uptake pathways likely varies and could contribute to differences in immune detection(57). For example, the endosomes accessed by reovirus via different routes may contain a distinct level of protease activity. Endosomes with higher protease activity may allow enhanced degradation and exposure of genomic dsRNA. Similarly, such endosomes may differ in the availability of yet to be identified cofactors required for entry or restriction factors that prevent entry. A possible proviral host factor could be the lipid composition of endosomes. It is known that host lipids impact the capacity of particles to undergo conformational transitions, generate pores, and escape endosomes(58).

The mechanism by which related dsRNA viruses, rotavirus and bluetongue virus, are sensed is not understood. Similar to reovirus, the RLR-MAVS signaling pathway is important for triggering an innate immune response against these viruses(59–68). Furthermore, viral replication is also likely dispensable for triggering an innate immune response to these viruses(63–68). Thus, our work highlights determinants that control the exposure and detection of genomic dsRNA of incoming reovirus particles. This study provides new information to understand innate immune recognition of multi-shelled dsRNA viruses that complete their replication cycle without the release of their PAMP from the particle.

